# Transcriptomic response to ISAV infection in the gills, head kidney and spleen of resistant and susceptible Atlantic salmon

**DOI:** 10.1101/2022.03.21.485193

**Authors:** O. Gervais, A. Papadopoulou, R. Gratacap, B. Hillestad, A.E. Tinch, S.A.M. Martin, R.D. Houston, D. Robledo

## Abstract

**Background:** Infectious Salmonid Anaemia virus (ISAV) is an orthomyxovirus responsible of large losses in Atlantic salmon (*Salmo salar*) aquaculture. Current available treatments and vaccines are not fully effective, and therefore selective breeding to produce ISAV-resistant strains of Atlantic salmon (*Salmo salar*) is a high priority for the industry. Genomic selection and potentially genome editing can be applied to enhance the disease resistance of aquaculture stocks, and both approaches can benefit from increased knowledge on the genomic mechanisms of resistance to ISAV. To improve our understanding of the mechanisms underlying resistance to ISAV in Atlantic salmon we performed a transcriptomic study in ISAV-infected salmon with contrasting levels of resistance to this virus.

**Results:** Three different tissues (gills, head kidney and spleen) were collected on 12 resistant and 12 susceptible fish at three timepoints (pre-challenge, 7 and 14 days post infection) and RNA sequenced. The transcriptomes of Infected and non-infected fish and of resistant and susceptible fish were compared at each timepoint. The results show that the responses to ISAV are organ-specific; an important response to the infection was observed in the head kidney, with up-regulation of immune processes such as interferon and NLR pathways, while in gills and spleen the response was more moderate. In addition to immune related genes our results suggest that other processes such as ubiquitination or ribosomal processing are important during early infection to ISAV. Moreover, the comparison between resistant and susceptible have also highlighted some interesting genes related to ubiquitination, intracellular transport or the inflammasome.

**Conclusions:** Atlantic salmon infection by ISAV revealed an organ-specific response, implying differential function during the infection. An early immune response was observed in the head kidney, while gills and spleen showed modest responses in comparison. Comparison between resistance and susceptible samples have highlighted genes of interest for further studies, for instance those related to ubiquitination or the inflammasome.

## BACKGROUND

Atlantic salmon (*Salmo salar*) is a valuable fish species farmed in several countries worldwide, and plays a major role supporting the economies of many rural communities. However, the sustainability of the industry is currently threatened by infectious diseases, which cause major economic losses.

One of the most threatening diseases for salmon farming is infectious salmon anaemia (ISA), caused by infectious salmon anaemia virus (ISAV) [1]. ISAV is listed by the OIE [2] as a notifiable disease and classified as a list II disease by the European Union fish health directive. This implies an active surveillance of the presence of the virus in fish farms and culling of stocks upon detection to avoid the transfer of the virus to other farms. Nonetheless, ISAV outbreaks have occurred in many salmon-producing countries, with the 2009 outbreak in Chile being particularly devastating, causing production losses of 75% [3–7]. ISAV belongs to the Orthomyxoviridae family and therefore is related to Influenza viruses [8]. The entry port of ISAV seems to be multiple; the gills are the main tissue of entry, but infection through the skin and pectoral fin is also possible [9, 10]. In Atlantic salmon this virus causes severe anaemia and haemorrhages, result of damage to the endothelial cells in peripheral blood vessels of all organs, which eventually leads to the death of the animal [3].

Nowadays, control of the disease mainly relies on farm surveillance and restriction of fish movements in infected/suspected farms. Some vaccines against ISAV have been developed, and they are extensively used in affected countries, however they do not confer full protection against the disease and therefore affected farms still have to isolate and cull their fish. A potential alternative is to produce stocks that are resistant to ISAV, either through selective breeding or genome engineering. Understanding molecular pathway and discovering functional genes involved in resistance / susceptibility to ISAV can significantly contribute to genomic selection and it is a necessary step to identify suitable targets for genome editing [11].

Previous *in vivo* studies on ISAV infection in Atlantic salmon have identified host genes potentially associated to resistance, such as hivep2 or TRIM25 [12, 13]. However, resistance to diseases tends to be multifactorial in nature, involving different biological pathways and complex organism-level responses that determine the balance in the host-pathogen relationship. In a previous study, we studied the response of Atlantic salmon to ISAV infection in the heart of resistant and susceptible fish [13]. To have a better vision of the fish systemic response during ISAV infection we have expanded our RNA sequencing study to three additional tissues: gills, head kidney and spleen. These tissues were selected due to their role in ISAV infection; head kidney and spleen are the main fish immune organs, while the gill is a key immune barrier and the main point of entry of ISAV. The transcriptomic response of these Atlantic salmon tissues to infection was assessed, and genetically resistant and susceptible animals were compared to better understand the genomic basis of resistance to ISAV.

## RESULTS

A total of 24 head kidney, 24 spleen and 24 gill samples were RNA sequenced (3’mRNA tag libraries), producing an average of 13M reads per sample. Principal component analyses showed a clear clustering of the samples of each tissue, but within each tissue no separation was observed between control and infected samples (Fig. 1A).

**Figure 1:**
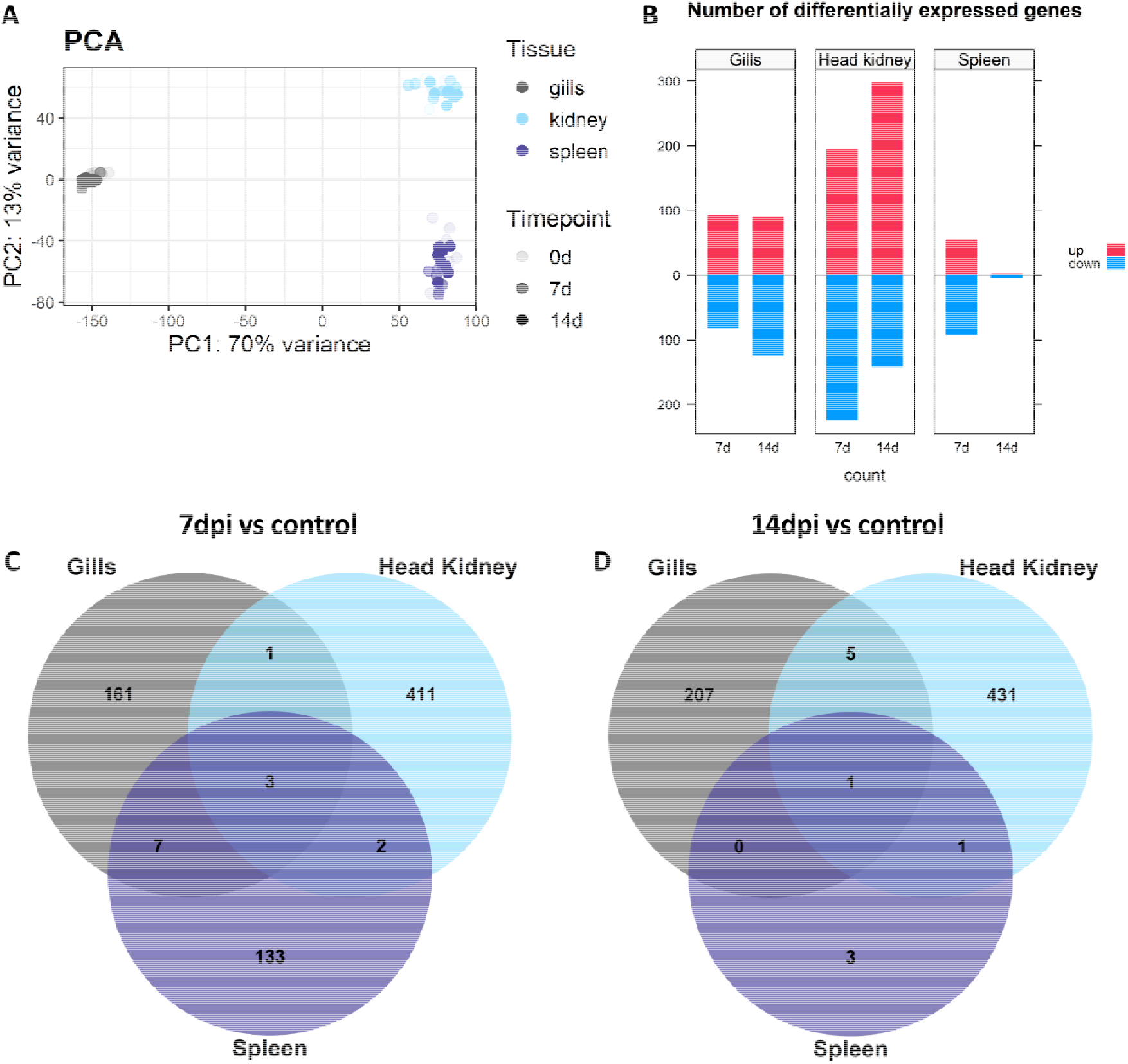
Differential gene expression between ISAV-infected and control fish. A) Principal Components Analysis showing the clustering of RNA-seq data; B) Diverted stacked bar chart showing differentially expressed genes (padj < 0.05) between control and infected samples in gill, head kidney and spleen, with up-regulated genes in red and down-regulated genes in blue; C) Venn diagram depicting the number of common and unique genes showing differential expression in each tissue at 7 dpi and D) 14 dpi compared to control.

### Differential expression analysis

Differential expression analysis between control and infected samples revealed 172, 417 and 145 genes differentially expressed for gills, head kidney and spleen respectively at 7 dpi (Fig. 1B). At 14 dpi, the number of differentially expressed genes is similar for gills and head kidney with 213 and 438 genes respectively, however in spleen only 4 genes were differentially expressed (Fig. 1B). Generally, a similar number of up- and down-regulated genes were observed in each comparison, except for head kidney 14 days post infection where a larger number of up-regulated genes were observed (Fig. 1B).

The differentially expressed genes are mostly organ-specific, however a small number of differentially expressed genes are common across the three tissues (Fig. 1C-D). There are 3 common genes at 7 dpi (Peptidyl-prolyl cis-trans isomerase FKBP5, FKBP5; Diencephalon/mesencephalon homeobox protein 1-B, DMBX1 and Serine protease 23) and just 1 at 14 dpi (Ankyrin repeat domain-containing protein SOWAHC, SOWAHC). Both FKBP5, an immunophilin, and DMBX1, a transcriptional repressor, were up-regulated at 7 days post infection (Fig. 2A-B), while Serine protease 23 is down-regulated at 7 days post infection (Fig. 2C). Finally, SOWAHC is part of the ankyrin repeat domain (ANKRD) family which mediates protein interactions and is down-regulated 14 days post infection (Fig. 2D).

**Figure 2:**
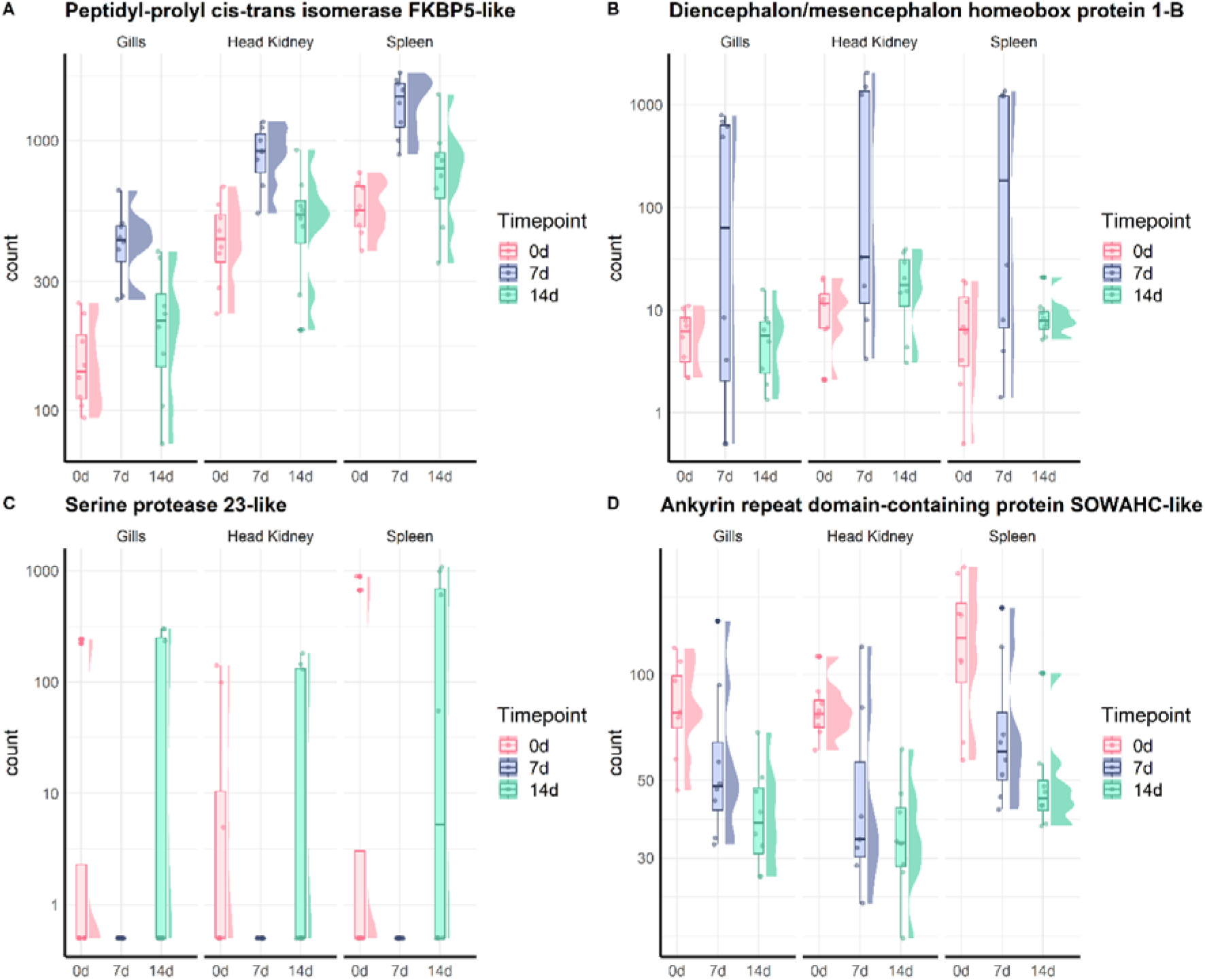
Gene expression patterns of common differentially expressed genes. Graph showing the number of normalised counts of common differentially expressed genes in gills, head kidney and spleen at 0, 7 and 14 dpi. Gene expression in each sample is represented with dots, and the distribution of the expression in each group is shown with a boxplot and a half-eye plot. A) Peptidyl-prolyl cis-trans isomerase FKBP5-like, B) Diencephalon/mesencephalon homeobox protein 1-B, C) Serine protease 23-like, D) Ankyrin repeat domain-containing protein SOWAHC-like.

### Response to ISAV in Atlantic salmon gills

In the gills, 172 and 213 differentially expressed genes were observed at 7 and 14 dpi respectively, with 46 of them shared between the two conditions (Fig. 3A). Based on gene ontology (GO) various biological processes (BP) were identified as up-regulated and, mostly, down-regulated at 7 and 14 days post infection (Fig 3B-C). At 7dpi, processes such as “response to stress”, “protein folding”, “metabolic process”, “immune system process”, “cell cycle” or “autophagy” are down-regulated, while 14 days after the infection “response to stress” and “immune system process” were slightly up-regulated, suggesting a late response to the infection in the gills, and others such as “ribosome biogenesis” and “ribonucleoprotein complex assembly” were down-regulated. Among the immune genes up-regulated at 14 dpi there are three genes related to major histocompatibility complex II and two C-C motif chemokines (Supplementary file1).

**Figure 3:**
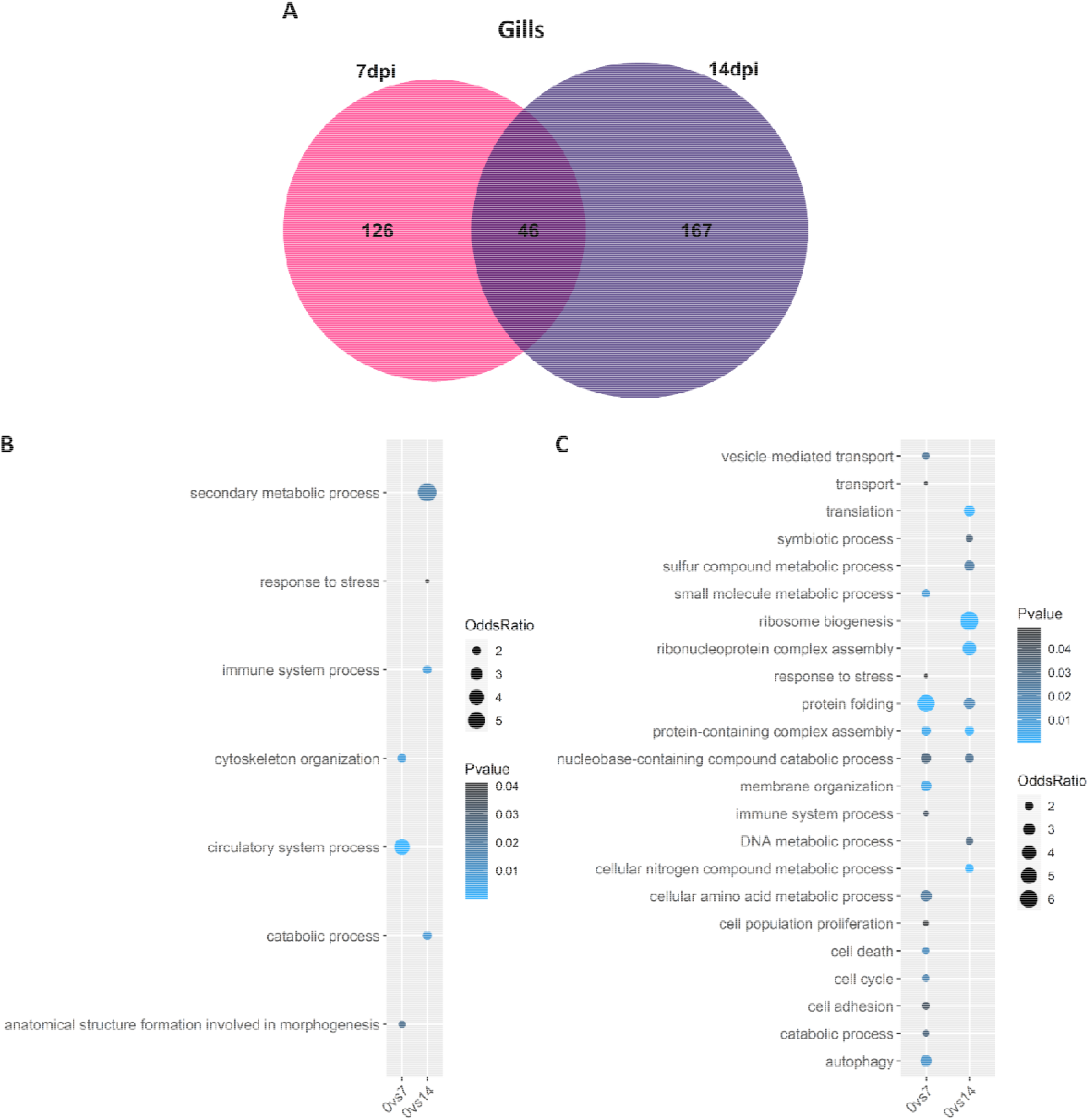
Common differentially expressed genes between 7 and 14 dpi in gills. A) Venn diagram depicting the number of common and unique genes showing differential expression at 7 and 14 dpi compared to control in gills. B-C) Bubblecharts showing enriched gene ontology in up-regulated (B) and down-regulated (C) genes at 7 and 14 days post infection compared to controls in gills.

### Response to ISAV in Atlantic salmon head kidney

In head kidney, 417 and 438 differentially expressed genes were found at 7 and 14 dpi respectively compared to control fish, with 126 in common between both timepoints (Fig. 4A). Many Biological Processes were enriched in the head kidney at both 7 (20 up- and 24 down-regulated) and 14 (14 up- and 6 down-regulated) days after the infection (Fig 4B-C). At 7 dpi the most up-regulated processes include “translation”, “ribosome biogenesis”, “protein targeting” and “catabolic process”, while other interesting terms such as “ribonucleoprotein complex assembly”, “cell death” or “cell cycle” show more moderate up-regulation. Similar to the gill results, down-regulated processes are “response to stress”, “cell division” and “cell cycle”. At 14 days post infection there were less enriched terms, but for example “ribosome biogenesis” and “ribonucleoprotein complex assembly” were up-regulated. Curiously, at both timepoints the cellular component term “ribosome” was up-regulated.

**Figure 4:**
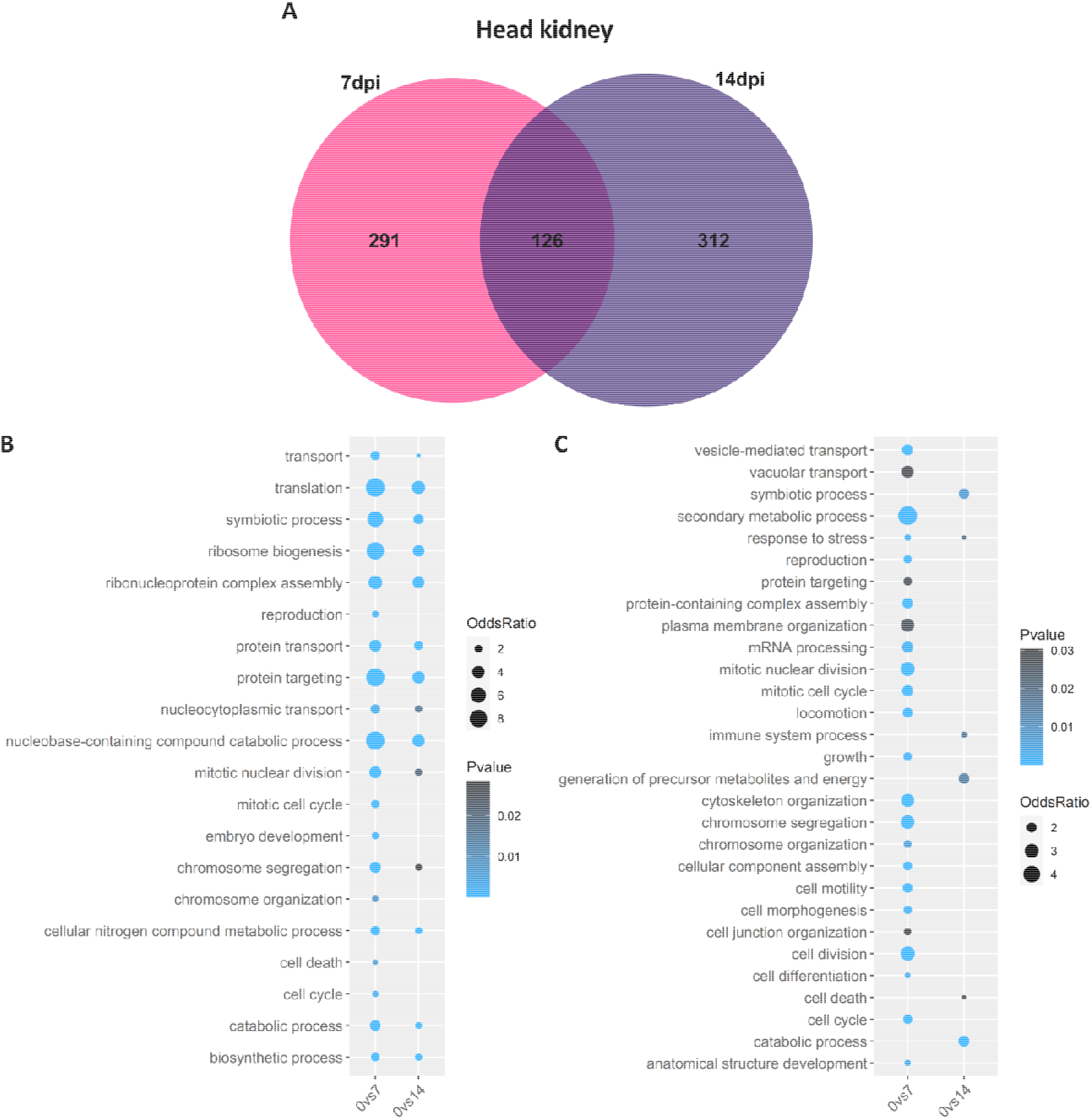
Common differentially expressed genes between 7 and 14 dpi in spleen. A) Venn diagram depicting the number of common and unique genes showing differential expression at 7 and 14 dpi compared to control in head kidney. B-C) Bubblecharts showing enriched gene ontology in up-regulated (B) and down-regulated (C) genes at 7 and 14 days post infection compared to controls in head kidney.

While we did not observe up-regulation of biological process related to immunity, we did observe up-regulation of multiple interferon related genes at both 7 (interferon alpha/beta receptor 1a, logFC= 0.99) and 14 days post infection (interferon-induced GTP-binding protein Mx, logFC= 3.59; interferon-induced protein 44, logFC= 2.01; interferon regulatory factor 7, logFC= 1.63) (Supplementary file2). Genes related to the NLR pathway such as proteins NLRC5 (logFC= 1.46) or protein NLRC3 (logFC= 0.95) were also up-regulated at 14 dpi.

### Response to ISAV in Atlantic salmon spleen

In the spleen only 145 and 5 genes were differentially expressed at 7dpi and 14dpi respectively, with no common genes between both conditions. Up-regulated biological processes in the spleen do not show an obvious connection to viral infection (e.g. “nitrogen cycle metabolic process” or “cell adhesion”; Fig. 5A-B). On the other hand, the down-regulated terms at 7 dpi include “response to stress”, “protein folding”, “cell death” and “cell cycle” (Supplementary file 3). Apparently, the spleen does not show a marked immune response to ISAV during the first stage of the infection.

**Figure 5:**
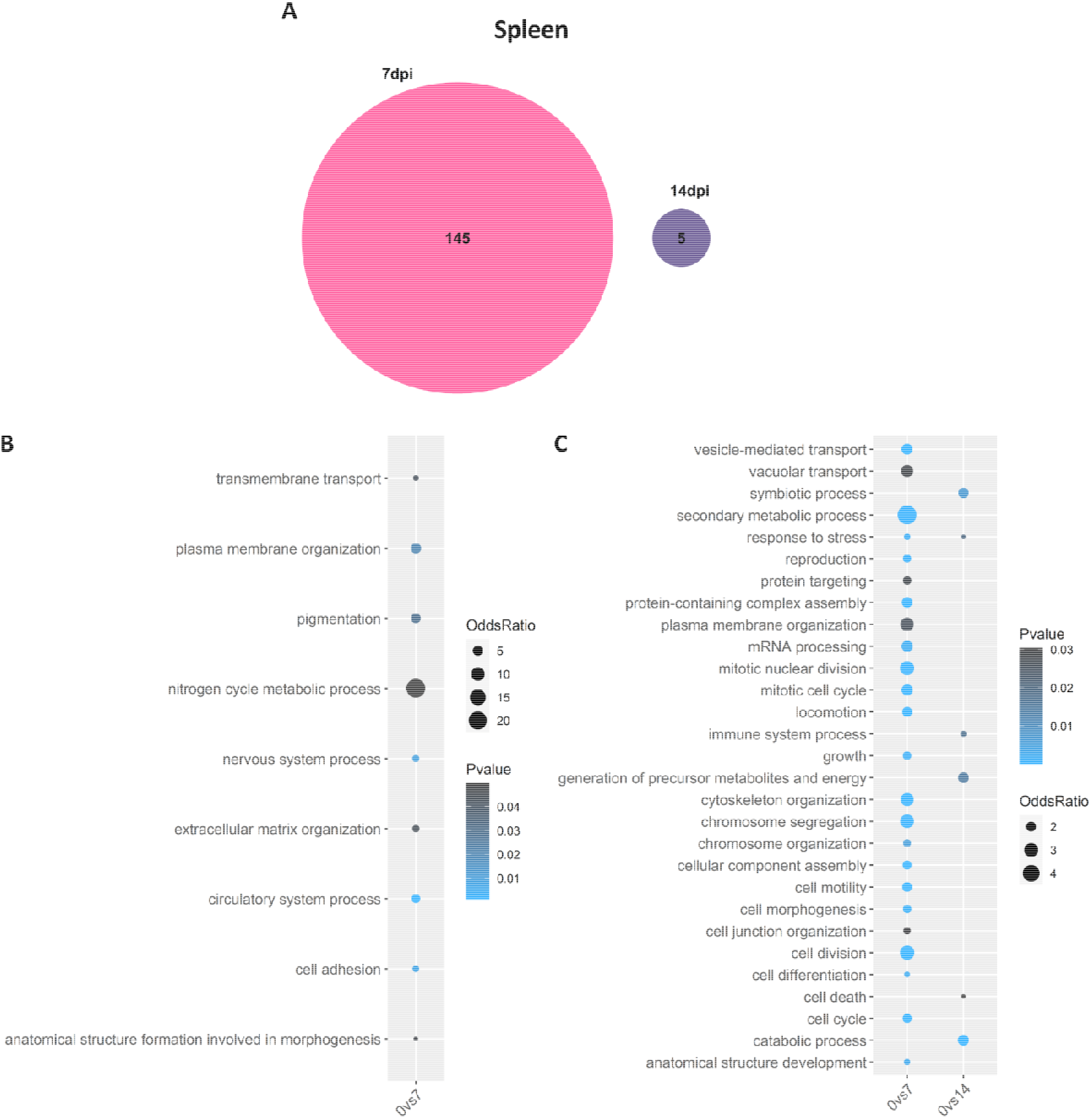
Common differentially expressed genes between 7 and 14 dpi in spleen. A) Venn diagram depicting the number of common and unique genes showing differential expression at 7 and 14 dpi compared to control in spleen. B-C) Bubblecharts showing enriched gene ontology in up-regulated (B) and down-regulated (C) genes at 7 and 14 days post infection compared to controls in spleen.

### Genomic signatures of resistance to ISAV in gill, head kidney and spleen

The transcriptomes of resistant and susceptible fish were compared for each tissue and timepoint (4 resistant vs 4 susceptible fish).

A small number of differentially expressed genes between resistant and susceptible samples were found in the gills (8-17 DEG per timepoint, Fig. 6 and Supplementary file 4). Some of those genes are related to the immune response. For instance, NACHT, LRR and PYD domains-containing protein 1-like (NLRP1), a key component of the inflammasome, is more expressed in susceptible samples at 7 dpi (logFC = −2.8). Also at 7 dpi, phospholipase A1 member A-like isoform X3 (PLA1A), involved in type I IFN production [14], is more expressed in resistant samples (logFC = 5.7). At 14 dpi, some differentially expressed genes are involved in response infections, such as transcriptional regulator ATRX-like (ATRX, logFC = 6.4), which plays a role in the maintenance of herpes simplex virus heterochromatin [15, 16]; the viral heterochromatin is formed during the lytic infection, where nucleosomes are assembled on the viral DNA and act as a epigenetic barrier to viral gene expression [17–19]. Polyadenylate-binding protein 1-like (PABC1) is less expressed in resistant fish at 14 dpi (logFC = −0.9), the cellular distribution of this gene is altered in various viral infections [20]. A gene involved in ubiquitination, ubiquitin-conjugating enzyme E2 D2 (UBE2D2), is also less expressed in resistant fish (logFC = −0.95).

**Figure 6:**
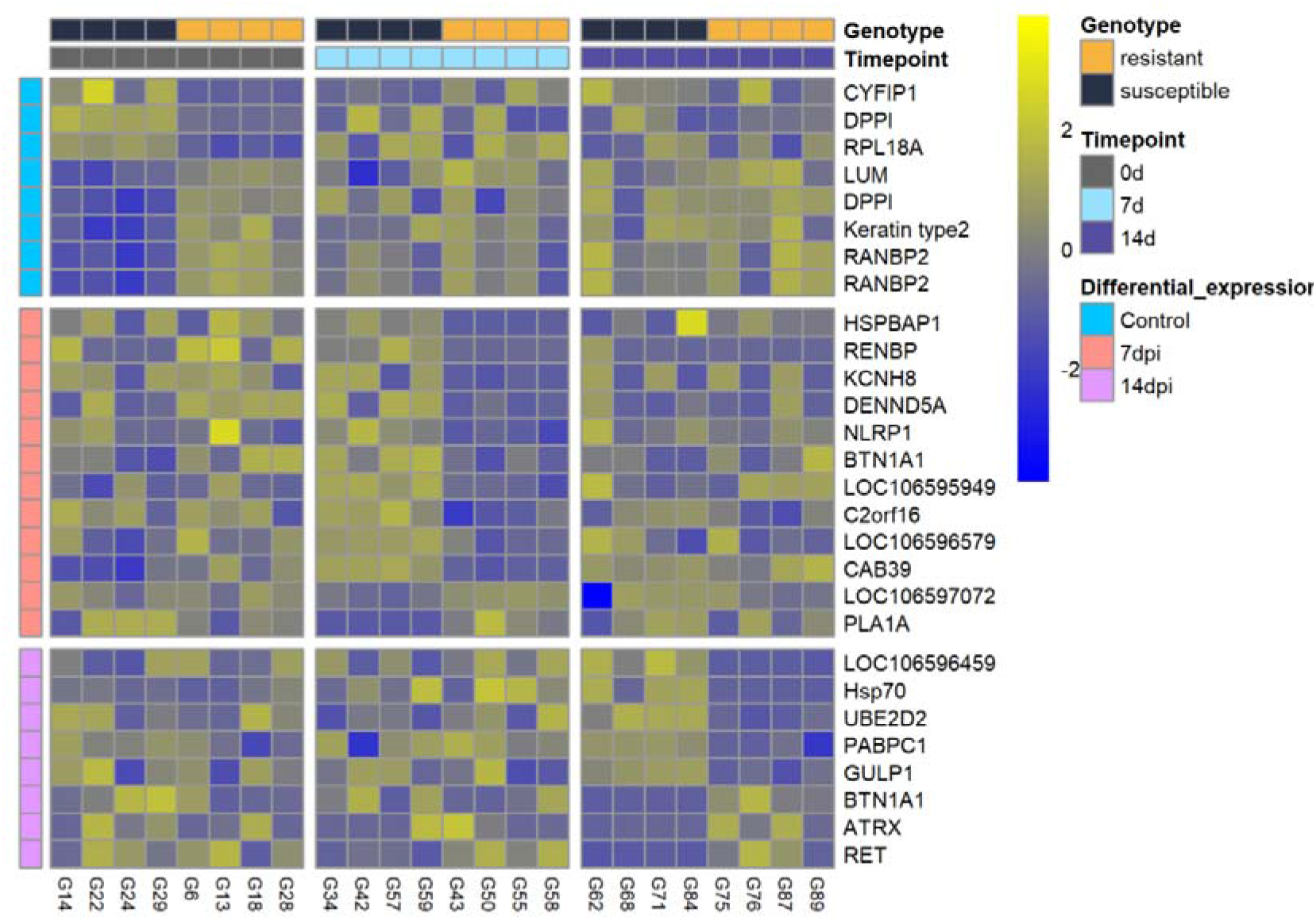
Heatmap showing the expression patterns of genes differentially expressed between resistant and susceptible fish in the gills at all three timepoints.

In head kidney, only 4 and 1 genes were differentially expressed between resistant and susceptible samples at 0 and 14 dpi, respectively. However, at 7 dpi a total of 152 genes were differentially expressed (Supplementary file 5). Interestingly, of those 152 genes only three genes were more expressed in resistant samples, with one of them being particularly interesting, nedd4-binding protein 2-like 1 a protein (N4BP2L1), which is involved in ubiquitination/neddylation [21, 22]. Most of the genes less expressed in resistant samples were involved in pathways related to the cytoskeleton or endosome (Supplementary file 7), both important cellular machineries used by virus for its intracellular transport.

Finally, in spleen a large number of genes were differentially expressed between resistant and susceptible samples at 0 dpi (264 genes), but not at 7 and 14 dpi (11-8 DEG) (Supplementary file 6). Those genes were mostly related to hemoglobin and ribosomes. At 7 dpi, terminal uridylyltransferase 7-like (TUT7), known to reduce Influenza A virus infection in early stages [23], was more expressed in resistant samples (logFC = 1.2). An inhibitor of phospholipase A2 (phospholipase A2 inhibitor 31 kDa subunit), a gene that can act as a regulator of the inflammation process [24–26], was significantly up-regulated in resistant fish at 14 dpi (logFC = 4.4).

## DISCUSSION

We have performed a multi-tissue RNA sequencing experiment to complement our previous work on the heart response to ISAV and gain a systemic view of the response of Atlantic salmon to this viral infection. ISA is a disease affecting the whole organism, where the virus circulates through the body of the fish using the blood vessels, multiplying in the epidermis of multiple tissues [27, 28]. In addition to the systemic immune response, each tissue can respond differently to the virus, and therefore a multi-tissue approach is important to understand this host-pathogen interaction. The three tissues studied in this experiment, gills, head kidney and spleen, where selected due to their involvement in the immune response and / or the entry of the virus into the fish.

The observed response to ISAV was markedly different in the three selected tissues, with a small number of common differentially expressed genes. Two of those genes could have an important role in the response to ISAV: FKBP5 and serine protease 23. Immunophilins such as FKBP5 have been previously reported to participate in viral replication during some infections such as HIV-1 [29]. Regarding serine protease 23, as an Orthomyxovirus ISAV possesses hemagglutinin (HA), a glycoprotein necessary for viral attachment and internalization into salmon cells [30, 31]. Influenza HA has to be cleaved for the virus to enter the cell, and that is done by a serine protease secreted by the host cell [32].

Our results show that head kidney displays a higher number of differentially expressed genes than gills and spleen. Head kidney is involved in haematopoiesis and cells found in this organ are capable of various immune functions, such as phagocytosis and antigen processing [33]. The increased expression of immune genes in head kidney, particularly the increased expression of interferon genes (irf2, irf4 and irf7) and the antiviral response triggered by the NLR pathway, reflect the immune function of this organ. A previous study showed similar results during early infection, with a more important up-regulation of interferon genes in the head kidney than in other tissues [34]. However, the interferon response was previously shown to be up-regulated also in other tissues, for instance in liver [35], and in these animals we previously found it down-regulated in the heart [13].

The difference observed during the infection between head kidney and the other two tissues is probably linked to their distinct immune function [36]. The spleen is the secondary lymphoid organ in teleost fish with an abundance of macrophages, responsible of erythrophagocytosis in early infection with ISAV [36, 37]. The increase of cell adhesion in spleen can potentially be related to the phagocytosis activity of macrophages as previously reported [37]. In gills, genes related to the major histocompatibility complex II (MHC II) and chemokine signalling were up-regulated. Previous studies during early ISAV infection have not shown an induction of MHC II, but up-regulation of MHC I has been reported [34]. Our results suggest that the gills not only act as the first barrier against ISAV, but that they are also capable of initiating specific immune responses against this pathogen.

In addition to immune related genes, other interesting regulatory pathways seem to play a role during early ISAV infection. Two of them are ubiquitination and neddylation, posttranslational modifications that modulate most cellular processes [38–40]. Ubiquitination-related genes were especially up-regulated in the head kidney of infected fish (7 at 7dpi and 15 at 14dpi). Moreover, two genes involved in this process were differentially expressed between resistant and susceptible fish; UBE2D2 was down-regulated in the gill of resistant fish, while N4BP2L1 was up-regulated in the head kidney of resistant fish. N4BP2L1 is involved in neddylation, a process that has been connected to resistance to infectious pancreatic necrosis virus (IPNV) in Atlantic salmon [41]. Nedd4 was also found to promote Influenza virus infection [42]. Additionally, some viruses need to hijack the host ubiquitination process for their own advantage [43, 44], and in fact the infection cycle of the orthomyxovirus Influenza requires ubiquitination for both cellular entry and replication [45]. Moreover, a previous study has highlighted the interaction of the s8ORF2 protein of ISAV with ubiquitin and interferon stimulated gene 15 (an ubiquitin-like protein) in Atlantic salmon cell culture (ASK) [46]. However, the molecular mechanisms underlying these interactions are still not known. In head kidney, two copies of the E3 ubiquitin-protein ligase HERC3 were upregulated in response to the virus at 14 dpi. In our previous study in the same population, a gene of the same family (HERC4) co-located with a putative QTL for resistance to ISAV [13]. Additionally, another gene of this family, HERC5, was previously described as an antiviral protein in Influenza virus infection, catalysing ISGylation of NS1 and avoiding its interaction with the antiviral protein kinase R (PKR), which reduces viral propagation [47]. Posttranscriptional modifications seem to play an important role during ISAV infection and it would be interesting to further investigate their role.

Our results also show an up-regulation of ribosomal protein genes in head kidney at both 7 and 14 dpi (e.g. RPS10, RPLP0, RPL15, RPL17 or RPL7). The role of ribosomal proteins (RPs) during viral infection has been investigated in multiple viruses, and interactions between RPs and viral proteins have been described in connection with viral protein biosynthesis as part of the normal replication cycle of the virus [48]. Different viruses prioritise certain RPs to complete their viral cycle. For example, in HIV-1 and WSSV (white spot syndrome virus) viral proteins interact with the ribosomal protein RPL7, while for RPS27a an interaction with a protein of Epstein-Barr viruses (EBV) has been described [48]. Additionally, many host proteins also interact with the viral ribonucleoprotein complex (RNP) of influenza virus, responsible for viral transcription and replication, and are fundamental for its transport and assembly [49, 50]. In our study, the ribonucleoprotein assembly complex process was up-regulated in head kidney in response to infection at both timepoints, potentially reflecting the hijack of the host machinery by ISAV as part of its infective process.

The comparison of susceptible and resistant fish highlighted certain genes of potential interest for further investigation, in addition to the previously mentioned genes involved in ubiquitination. The largest differences were observed in the head kidney, where interestingly the difference between resistant and susceptible fish does not stem from differences in immune pathways, but mostly a down-regulation of various pathways involved in intracellular transport: cytoskeleton, microtubules and endosomes. Many viruses exploit these cellular processes for cell entry and intracellular transport [51, 52], and endosomes and lysosomes have been previously reported to be the entry way of ISAV into the cell [53]. Their down-regulation in resistant animals may affect viral replication by reducing viral entry and trafficking on infected cells, but it is also possible that susceptible animals simply have a higher expression of these pathways as a consequence of a more severe viral infection. These processes and associated genes require more investigation to validate their role in resistance / susceptibility to ISAV.

In the gills of resistant fish, we observed an increase of the expression of NLRP1 at 7 dpi, a core protein of the inflammasome. Moreover, two other genes involved in the inflammasome were modulated in response to ISAV. Interleukin 1 was down-regulated in gills and caspase 1 up-regulated in head kidney at 7 and 14 dpi respectively when compared to controls. The inflammasome is a key regulator of the host response against pathogens, which can promote cell death to clear infected cells [54, 55]. There are multiple types of inflammasomes (NLRP3, NLRP1, AIM2, NAIP-NLRC4, etc.) which are activated via different pathways, for example the NLRP3 inflammasome is activated by several viral viroporins [56]. Inflammasomes are highly regulated since inappropriate or excessive activation can lead to significant pathology [57]. Further, some viruses such as orthopoxvirus and Influenza virus can inhibit inflammasome signalling [55]. Inflammasomes are understudied in fish and it would be interesting to investigate their role during ISAV infection.

In the spleen, two genes up-regulated in resistant fish seem to be interesting for ISAV resistance. The first one is TUT7, a potent antiviral factor during early stages of RNA virus infection, and its deletion leads to increased IAV and orsay virus mRNA [23]. The other one is Phospholipase A2 inhibitor (PLA2); two inhibitors of phospholipase A2 were previously found to be up-regulated in infected fish showing delayed mortality, but not in early mortalities [25]. Additionally, flavivirus West Nile virus was found to manipulate lipid homeostasis using PLA2 to facilitate its replication [26].

## CONCLUSIONS

The transcriptomic analysis of ISAV-infected Atlantic salmon has revealed a complex tissue-specific response. Each tissue responds differently to the infection with the head kidney presenting a high immune response compared to gills and spleen. Comparison of genetically resistant and susceptible animals suggests there is not a single clear resistance mechanism, which is consistent with the polygenic nature of ISAV resistance in Atlantic salmon. Our results also reveal that resistance to ISAV may not only be dependant of purely immune pathways and cellular mechanisms such as posttranslational modification or various intracellular transport pathways may also contribute to ISAV resistance. Further validation through functional studies are necessary to explore the importance of these genes and pathways, and reveal the cellular mechanisms underlying resistance to ISAV in Atlantic salmon.

## METHODS

### Disease challenge and sampling

The population used for the ISAV challenge experiment comprised 2,833 parr Atlantic salmon (mean weight 37.5 ± 9.2 g) from 194 nuclear families originating from Benchmark Genetics breeding programme. The challenge experiment and sampling were conducted in the facilities of VESO Vikan (Norway). All the disease challenge and sampling protocol was previously described in detail in [13]. After acclimatation of the fish during three week, 300 carrier fish (Atlantic salmon from the same population) were intraperitoneally injected with 0.1 mL of ISAV (Glaesvær, 080411, grown in ASK-cells, 2 passage, estimated titre 10^6^ PFU / mL [58]) and introduced to the challenge tank with naïve fish. Fish and tanks were monitored on daily basis, mortalities were registered and sampled, environmental parameters were also recorded. The trials ends when the mortality reach the level near zero. In addition, gills, head kidney and spleen of 30 challenge fish were collected at three timepoint (pre-infection, 7 dpi and 14 dpi) into TRI Reagent (Sigma, UK) and stored at −80°C until RNA extraction.

### RNA extraction and RNA sequencing

For each timepoint (control, 7 dpi and 14 dpi) 4 fish with high breeding values for resistance and 4 fish with low breeding values for resistance, representing 8 different families, were selected according to their estimated genomic breeding value for ISAV resistance [13]. Gills, head kidney and spleen RNA samples from the same fish were extracted from preserved tissue samples in TRI reagent (Sigma, UK) and RNA extracted following the manufacturer’s instructions (n= 24 per tissue; control = 8; 7 dpi = 8; 14 dpi = 8). The RNA pellet was eluted in 15 μL of nuclease-free water and quantified on a Nanodrop 1000 spectrophotometer (NanoDrop Technologies) prior to DNAse treatment with QuantiTect^®^ Reverse Transcription kit (Qiagen). The quality of the RNA was examined by electrophoresis on a 1% agarose gel (Sigma Aldrich), prepared in Tris-Acetate-EDTA (TAE) buffer, stained with 1% SYBR Safe (Sigma Aldrich) and run at 80 V for 30 min. Sample concentration was measured with Invitrogen Qubit 3.0 Fluorometer using the Qubit RNA HS Assay Kit (ThermoFisher Scientific). The 3’mRNA tag-seq libraries were prepared by Oxford Genomic Centre, and sequenced on a Illumina Novaseq6000 with an average of 13.1M reads (minimum 9.3M).

### RNA-Seq analyses

Raw reads were quality trimmed using Trimgalore v0.6.3. Briefly, adapter sequences were removed, low quality bases were filtered (Phred score < 20) and reads with less than 20 bp were discarded. Trimmed reads were pseudo aligned against the Atlantic salmon reference transcriptome (ICSASG_v2 Annotation Release 100; Lien et al., 2016) using kallisto v0.44.0 [60]. Transcript level expression was imported into R v4.0.2 [61] and summarised to the gene level using the R/tximport v1.10.1 [62]. Differential expression analysis was performed using R/Deseq2 v1.28.1 [63], and genes with False Discovery Rate adjusted p-values < 0.05 were considered to be differentially expressed. Gene Ontology (GO) enrichment analyses were performed in R v.3.5.2 using Bioconductor packages GOstats v.2.54.0 [64] and GSEABasse v.1.50.1 [65]. GO term annotation for the Atlantic salmon transcriptome was obtained using the R package Ssa.RefSeq.db v1.3 (https://gitlab.com/cigene/R/Ssa.RefSeq.db). The over-representation of GO terms in differentially expressed gene lists compared to the corresponding transcriptomes (gills, head kidney or spleen) was assented with a hypergeometric test. A GO terms was considered enriched if it showed ≥5 DE genes assigned and a p-value <0.05.

### Ethics statement

The challenge experiment was performed at VESO Vikan with approval from the Norwegian Food Safety Authority, National Assignments Department, approval no.16421, in accordance with the Norwegian Animal Welfare Act.

### Data availability

RNA sequencing raw reads have been deposited in the NCBI’s Short Read Archive (SRS) repository with accession number PRJNA780199.

## DECLARATION

### Ethics approval and consent to participate

The challenge experiment was performed at VESO Vikan with approval from the Norwegian Food Safety Authority, National Assignments Department, approval no. 16421, in accordance with the Norwegian Animal Welfare Act.

### Consent for publication

Not applicable

### Author contributions

RH, SAM and DR were responsible for the concept and design of this work. BH and AE were responsible for the disease challenge. OG, AB, AP and RG analysed the data. OG, DR and RH drafted the manuscript. All authors reviewed and approved the manuscript.

## Acknowledgements

Not applicable

